# Proteomic Profiling of Enriched Nuclei Provides a Nuclear Proteome Resource and Protein Interaction Landscape for *Trypanosoma cruzi*

**DOI:** 10.64898/2026.08.25.746806

**Authors:** Rafael Fogaça de Almeida, Matheus Fernandes, Lyris Martins Franco de Godoy

## Abstract

The processes such as DNA replication, transcription, and repair are often modulated by specific nuclear proteins, protein–protein interactions (PPIs), and post-translational modifications (PTMs). In *Trypanosoma cruzi*, however, the nuclear proteome and interactome have not been systematically mapped, limiting the interpretation of nuclear regulatory processes. Here, we report a nuclear proteome resource generated from intact nuclei isolated from *T. cruzi* and analyzed by high-resolution Orbitrap LC–MS/MS, integrating proteome profiling, computational interaction network inference, and exploratory crosslinking mass spectrometry (XL-MS). Proteome profiling identified 1,734 proteins in the nuclear fraction, including 316 proteins identified with PTM-containing peptides. Subcellular localization prediction and Gene Ontology analysis support nuclear enrichment and highlight functions related to transcription, RNA metabolism, and genome maintenance. The *in silico* interaction network derived from STRINGDB organizes the proteins into functional clusters, including a histone-associated interaction neighborhood. In parallel, XL-MS identified 26 residue-resolved interprotein crosslinks involving 36 proteins and detected PTMs at or near linked residues. Together, these data support reuse for comparative nuclear proteomics, multiomics integration, and prioritization of candidates for future functional studies.

## 1 Introduction

The cell nucleus hosts essential processes such as DNA replication, transcription, and repair, which are required for survival, growth, and differentiation. These processes are carried out by proteins that act individually or recruit multiprotein complexes to perform their function. Protein-protein interactions (PPIs) can be stable or transient and are frequently regulated by posttranslational modifications (PTMs). Histones, in particular, carry a wide variety of PTMs that can influence transcription and chromatin organization by directly altering nucleosome dynamics or by creating binding sites for effector proteins and complexes involved in DNA repair and other essential nuclear functions [1].

Crosslinking mass spectrometry (XL-MS) is a powerful methodology to capture native PPIs in cells and organelles and has been applied to characterize interactomes in mammalian systems, including the human cell nucleus [2,3]. In addition to providing residue-level distance restraints, XL-MS allows the identification of PTMs that are close to, and possibly involved in, the interaction sites [2]. Chemical crosslinking can also stabilize endogenous complexes during sample handling, helping to preserve labile or transient interactions and reducing post-lysis rearrangements [4].

*Trypanosoma cruzi*, the etiologic agent of Chagas disease [5], is an interesting model for studying nuclear PPIs. Despite having predominantly post-transcriptional gene expression regulation, we recently showed that *T. cruzi* displays a broad repertoire of histone PTMs (hPTMs), similar to those described for humans and other eukaryotes [6,7], raising questions about how they contribute to processes such as DNA repair and broader epigenetic regulation. Expanding and annotating the *T. cruzi* nuclear proteome, as well as mapping the nuclear protein-protein interactions, particularly those involving histones, are therefore important steps toward understanding how replication, transcription, DNA repair, and other essential functions are organized in this parasite. In this context, we isolated intact nuclei and conducted integrative proteomic and computational analyses to generate a nuclear proteome and interactome resource for *T. cruzi*.

We provide a high-quality nuclear proteome from epimastigotes with >1700 proteins, accompanied by subcellular localization, solubility/membrane predictions, and functional enrichment analyses. We further present *in silico* nuclear interaction networks predicted using the STRING database, complemented by exploratory XL-MS data that provide residue-level evidence for 26 interprotein crosslinks involving 36 proteins and PTM annotations at or near linked sites.

## 2 Materials and Methods

The overall workflow for the analysis of the *T. cruzi* nuclear proteome and PPIs is outlined in Figure 1, while the experimental procedures are described in detail below.

**Figure 1.**
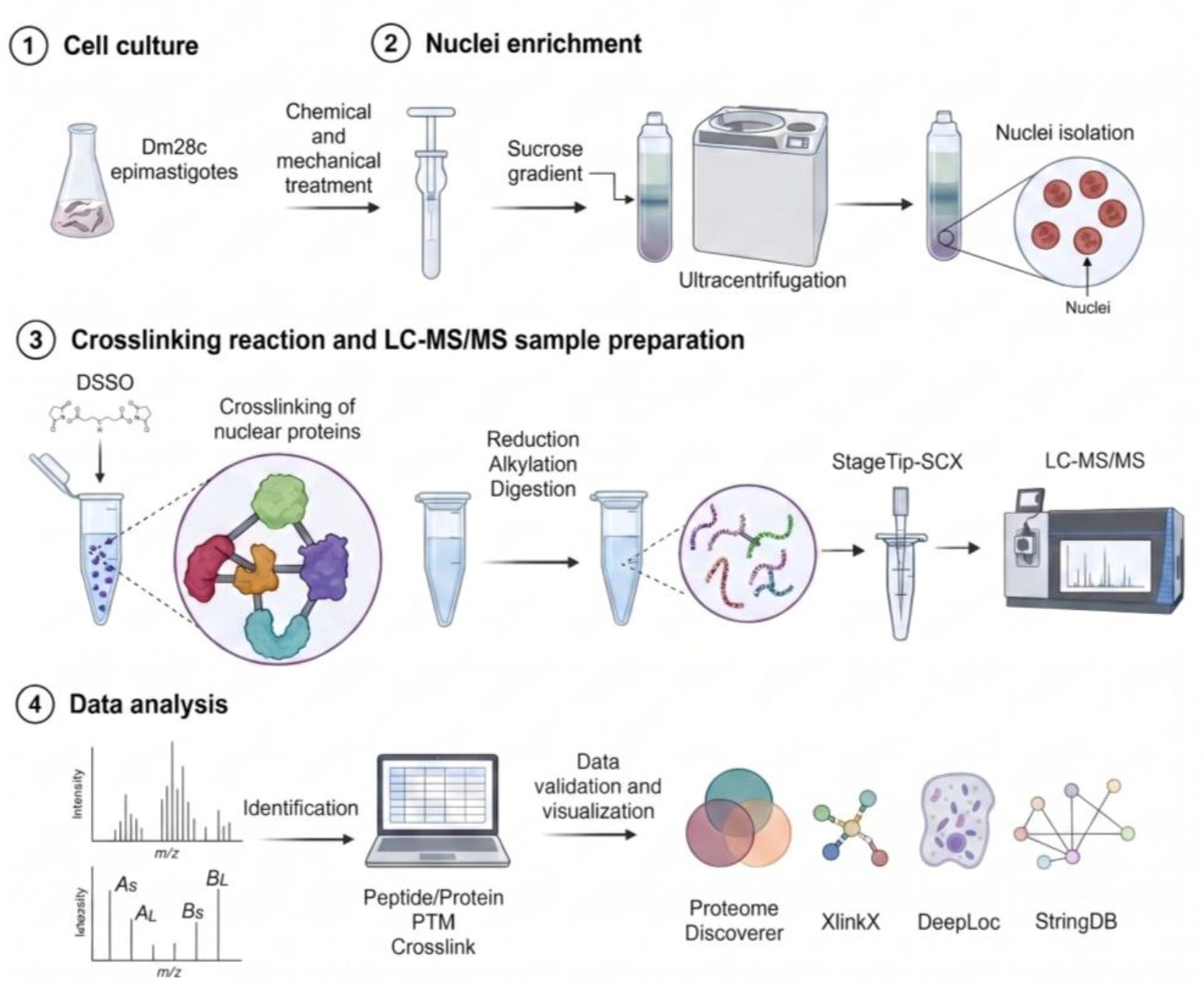
Integrated workflow for nuclear proteome and interactome analysis in *T. cruzi*. (1) *T. cruzi* Dm28c clone epimastigotes were harvested during exponential growth. (2) Cells were disrupted by chemical and mechanical lysis using a Dounce homogenizer, and intact nuclei were enriched by ultracentrifugation through a discontinuous sucrose gradient. (3) Purified intact nuclei were crosslinked with DSSO, followed by nuclear lysis, protein reduction and alkylation, and tryptic digestion. Peptides were prefractionated by strong cation exchange chromatography (SCX-StageTip) and analyzed by nanoLC-MS/MS. (4) Mass spectrometry data were processed using well-established proteomics platforms for peptide/protein identification, PTM annotation, and crosslinked peptide identification, and the results were subsequently analyzed and explored using different downstream computational tools for validation, statistical analysis, and network visualization.

### 2.1 Cell culture and intact nuclei enrichment

*T. cruzi* Dm28c epimastigotes in exponential growth phase were cultured in Liver Infusion Tryptose (LIT) medium [8] supplemented with 10% fetal bovine serum and incubated without agitation at 28 °C. *T. cruzi* nuclei were enriched based on the protocol previously described [9] with some modifications. Briefly, 1×10^9^ exponential-phase epimastigotes were resuspended in hypotonic buffer (10 mM Tris-HCl (pH 7.4), 10 mM NaCl, 1 mM MgCl_2_, 1 mM MnCl, 5 mM βmercaptoethanol), and cell turgidity was confirmed by optical microscopy. Nonidet P40 (0.5% v/v) and protease inhibitor (Complete Mini, Roche) were added, and the cells were lysed with a Dounce homogenizer. Cell lysis was checked by optical microscopy. Lysate osmolarity was restored by adding sucrose to a final concentration of 0.25 M, and the lysate was slowly layered onto 1 mL of 0.58 M sucrose in hypotonic buffer, followed by centrifugation at 2,000 x g for 10 min at 4 °C. The top layer (cytosolic fraction) was removed, and the pellet was resuspended in 1.9 M sucrose in hypotonic buffer. The resuspended pellet was carefully loaded into an ultracentrifuge tube (SW65) containing a discontinuous sucrose gradient (bottom to top: 2.30 M; 2.10 M; and 2.01 M sucrose, all prepared in hypotonic buffer with protease inhibitors). The tube was ultracentrifuged (141,000 x g; 3 h; 4 °C), and the pellet containing the enriched intact nuclei was stained with the blue-fluorescent 4’,6-diamidino-2-phenylindole (DAPI) and analyzed by fluorescence microscopy.

### 2.2 Protein crosslinking and proteolytic digestion

Crosslinking of nuclear proteins was based on the protocols previously described for human cells [2,10]. A single biological nuclear preparation was used for all experiments. The nuclei sample was washed with crosslinking reaction buffer (20 mM Hepes, 10 mM NaCl, 1 mM MgCl_2_6H2O, pH 8), and divided into four equal aliquots, each corresponding to one dissuccinimidyl sulfoxide (DSSO) concentration (0.2, 0.6, 1.0, or 1.5 mM). The DSSO concentration series (0.2–1.5 mM) was incorporated into the experimental design to sample a range of crosslinking conditions and increase the recovery of crosslinked peptides. All DSSO-treated samples were subsequently analyzed by LC–MS/MS and included in the final dataset; therefore, they represent analytical runs rather than a separate optimization or quality-control experiment. Crosslinking reactions were performed for 25 min at room temperature. The reaction was quenched with 50 mM Tris-HCl (pH 8) for 10 minutes. Nuclei were pelleted by centrifugation at 3,000 x g, 4°C for 10 minutes. Total nuclear protein extract was obtained by lysing nuclei in 50 mM Tris (pH 8) and 2% SDS, heating at 95 °C for 3 min, sonicating, and centrifuging at 20,000 x g for 30 min. Proteins were acetone-precipitated and pelleted at 6,000 x g for 15 min at 4 °C. The pellet was resuspended in 50 mM ammonium bicarbonate and 8 M urea, reduced with 15 mM dithiothreitol for 1 h at room temperature. Alkylated with 50 mM iodoacetamide for 2 h in the dark. Alkylation was quenched with thiourea at a final concentration of 150 mM, and the samples were diluted fourfold with 50 mM ammonium bicarbonate. Proteins were digested with trypsin (1:100, enzyme:protein) at 37 °C for 16 h. The peptides were desalted in C18-StageTips [11].

### 2.3 Peptide fractionation

Desalted peptides were fractionated on SCX-StageTip as previously described [10]. Briefly, homemade SCX-StageTips were prepared by packing three layers of EmporeTM sulfonic acid strong cation exchange disks into 200-µL pipette tips and peptides were eluted with buffers containing increasing ammonium acetate (AmAc) concentrations (200, 500, and 1,000 mM). The 200 mM and 500 mM eluates were combined. A total of 8 samples (four samples of different DSSO concentrations with two different AmAc fractions/combinations) were desalted on C18-StageTip, and analyzed by nanoLC–MS/MS in duplicate, totaling 16 .raw files. The relationship between DSSO concentrations, SCX fractions, LC–MS/MS runs, deposited files, and downstream analyses is summarized in Table S1.

### 2.4 LC-MS/MS analysis

The analyses for the proteome and interaction partners were performed using an Ultimate 3000 RSLCnano system coupled to an Orbitrap Fusion Lumos mass spectrometer (Thermo Scientific) at the Carlos Chagas Institute mass spectrometry platform (RPT02H). Peptide separation was performed on an in-house packed analytical C18 column (15 cm × 75 μm internal diameter) packed with 3 μm ReproSil-Pur C18 resin (Dr. Maisch GmbH, Germany). Peptides were separated over a 120 min gradient at a flow rate of 250 nL/min using mobile phase A consisting of 0.1% formic acid in water and mobile phase B consisting of 0.1% formic acid in 95% acetonitrile. The acquisition parameters were defined according to previously reported methodologies [10,12]. The acquisition method was CID-MS2-MS3-ETD-MS2. Data were acquired in positive ion mode. The ion source was operated with a spray voltage of 2.3 kV and an ion transfer tube temperature of 175 °C. Internal mass calibration was enabled using EASY-IC. Data were acquired in datadependent acquisition (DDA) mode using a duty cycle of 5 s. Full MS scans (MS1) were acquired in the Orbitrap analyzer at 60,000 resolution (at m/z 200) over an m/z range of 375–1500 using profile mode. MS1 scans were acquired with quadrupole isolation enabled, AGC target set to 4 × 10⁵ ions, automatic maximum injection time (up to 50 ms), and one microscan per spectrum. Only precursor ions with charge states between +2 and +8 were selected for fragmentation, while precursors with undetermined charge states were excluded. Dynamic exclusion was enabled to minimize repeated fragmentation of the same precursor ions. Precursors were excluded after one selection for 45 s using a ±10 ppm mass tolerance window. Isotope exclusion was enabled, and precursor selection required a minimum intensity threshold of 2.0 × 10⁴. MS2 spectra were acquired by CID fragmentation in the Orbitrap analyzer at 30,000 resolution using a quadrupole isolation window of 1.6 m/z, normalized collision energy of 25%, AGC target of 5 × 10⁴ ions, and maximum injection time of 70 ms. Spectra were acquired in profile mode with one microscan. MS2 precursor selection was restricted to charge states between +2 and +6. Targeted mass difference triggering was configured for DSSO cross-links using a characteristic doublet mass difference of Δ31.9721 Da. The method required detection of two precursor ions within a relative partner intensity range of 30–100% and within a tolerance of ±10 ppm. Triggered MS3 scans were acquired in the ion trap using rapid scan mode, an isolation window of 2 m/z, and CID fragmentation with 35% collision energy. MS3 scans were acquired with an AGC target of 2 × 10⁴ ions and a maximum injection time of 120 ms in centroid mode. In parallel, ETD-based MS2 scans were also acquired in the Orbitrap analyzer at 30,000 resolution over an m/z range of 150–2000. ETD scans employed a quadrupole isolation window of 1.6 m/z, calibrated charge-dependent ETD parameters, AGC target of 5 × 10⁴ ions, and a maximum injection time of 100 ms. ETD spectra were acquired in profile mode without supplemental activation.

### 2.5 Proteome and PTM searches

Raw LC–MS/MS data were processed in Proteome Discoverer v3.1 SP1 using the built-in workflow template PWF_Tribrid_SequestHT_MSAmanda_Percolator, which integrates Sequest [13] and MS Amanda [14] search engines, followed by Percolator [15] to assign statistical confidence metrics (q-values and posterior error probabilities) to peptide-spectrum matches (PSMs). Two independent database search types were performed on the same 16 .raw files: a standard proteome search (“Proteome”) using default parameters, and a second search specifically optimized for post-translational modification identification (“PTMs”). Both searches were performed against the *Trypanosoma cruzi* Dm28c 2017 protein sequence database downloaded on October 10, 2025, from TriTrypDB [16] v.68, supplemented with a common contaminants database. In each search engine, trypsin was specified as the proteolytic enzyme (full digestion), allowing up to four missed cleavages, and peptide and protein identifications were filtered at 1% false discovery rate (FDR) in Percolator. Precursor and fragment ion mass tolerances were set to 10 ppm and 0.02 Da, respectively, for both Sequest HT and MS Amanda 3.0 search engines. Carbamidomethylation of cysteine (+57.021 Da) was defined as a fixed modification for both searches, as well as methionine oxidation (+15.995 Da) as a variable modification. For the PTM search, the following dynamic modifications were included: lysine acetylation (+42.011 Da), monomethylation and dimethylation of lysine/arginine (+14.016 Da and +28.031 Da, respectively), trimethylation of lysine (+42.047 Da), and phosphorylation of serine, threonine, and tyrosine residues (+79.966 Da). N-terminal modifications including acetylation, methionine loss, and methionine loss followed by acetylation were also considered for both searches. MS/MS spectra were preprocessed using the Spectrum Selector node with isotope pattern precursor reevaluation enabled and precursor mass filtering between 350 and 5000 Da. Only full-scan MS/MS spectra were considered, using a minimum signal-to-noise threshold of 1.5. Identification validation was performed using the Percolator algorithm with a concatenated target-decoy strategy and q-value-based false discovery rate (FDR) estimation. Strict FDR thresholds were set to 1% at the PSM, peptide, and protein levels, while relaxed thresholds were set to 5%. Only high-confidence peptides were retained for downstream analyses. In the consensus workflow, results were globally merged by search engine type using the PSM Grouper node. Protein grouping was performed using the strict parsimony principle, and only master proteins were considered for modification and peptide position annotations. Modification site localization confidence was controlled using the ptmRS localization probability with a minimum site probability threshold of 75%; therefore, PTMs reported in this study correspond to confidently localized modification sites rather than peptide-level observations alone. PTM-containing peptide-spectrum matches were validated using Percolator at a strict 1% FDR (q-value-based) threshold. The proteins identified in “Proteome”, “PTMs” or both (“Proteome-PTMs”) searches were compiled in Table S2.

### 2.6 Protein subcellular annotation and functional enrichment

DeepLoc v.2.1 [17] was used to classify proteins by subcellular compartment, to annotate nuclear targeting features including predicted nuclear localization signals (NLS) and/or nuclear export signals (NES) – and to infer solubility versus membrane association, including membrane subtype assignments. Functional enrichment of Gene Ontology (GO) [18] slim terms across cellular component, biological process, and molecular function categories was performed using the TriTrypDB GO search tool [16], applying a p-value cutoff of 0.05 and multiple-testing corrections (Benjamini–Hochberg and Bonferroni).

### 2.7 In silico interactome analysis and network clustering

Proteins identified in the nuclear preparation were mapped in STRINGDB v.12 [19] using StringApp [20] and visualized in Cytoscape v.3.10.4 [21]. The released interaction resource was retrieved in full STRINGDB mode using *T. cruzi* as the species and applying a minimum required interaction score of 0.7. Network clusters were identified using the Markov Cluster Algorithm (MCL) [22] on the released network, with an inflation value of 4 (default). For figure visualization, edges were further restricted to a high-confidence subset (score ≥ 0.9) supported by experimental or curated database evidence. Clusters were subsequently analyzed for functional enrichment [20], and the most enriched GO Biological Process term (with FDR) was used to annotate clusters in downstream visualization.

### 2.8 XL-MS–derived interactome analysis and annotation

Crosslinked peptides were searched using XlinkX v.3.1 [12] implemented within Proteome Discoverer v.3.1 SP1, using the MS2-MS2-MS3 acquisition template. The *Trypanosoma cruzi* Dm28c 2017 database, downloaded on October 10, 2025, from TriTrypDB v.68, was used for searching. For peptide identification, spectra were searched with the Sequest HT search engine using trypsin as the proteolytic enzyme, allowing up to five missed cleavages. Precursor mass tolerance was set to 20 ppm for Sequest HT searches, while fragment mass tolerance was set to 0.6 Da. In the XlinkX search node, precursor mass tolerance was set to 10 ppm, FTMS fragment tolerance to 20 ppm, and ITMS fragment tolerance to 0.5 Da. Peptide lengths ranged from 5–150 amino acids, and up to 10 peptide candidates per spectrum were retained. Carbamidomethylation of cysteine (+57.021 Da) was set as a static modification. Variable modifications included methionine oxidation (+15.995 Da), protein N-terminal acetylation (+42.011 Da), DSSO hydrolyzed adducts on lysine residues (+176.014 Da), and DSSO Tris adducts on lysine residues (+279.078 Da). The DSSO cross-link modification (+158.004 Da on lysine residues) was specified in the XlinkX detection node, and protein N-terminal cross-linking was enabled. A maximum of four dynamic modifications per peptide and three equal modifications per peptide were allowed. Separate processing branches were applied for CID and ETD fragmentation spectra. False discovery rate (FDR) estimation was performed using a concatenated target-decoy strategy. Peptide-spectrum matches (PSMs), peptides, and cross-links were filtered to a strict FDR threshold of 1%. FDR control of crosslinking identification was applied in two steps: (i) in XlinkX Validator, which validates identifications at the spectrum level (MS2 and MS3) and classifies inter- and intra-links, and (ii) in the consensus workflow, controlling the false positive rate at the crosslink spectrum match (CSM) level [23]. Inter- and intra-protein cross-links were evaluated separately. Protein grouping was performed according to the strict parsimony principle. In XlinkX Search, dynamic modifications were enabled to annotate PTMs on crosslinked peptides, using the most frequent histone-associated PTMs previously reported in *T. cruzi* [6]: acetylation (+42.01 Da, K), methylation (+14.01 Da, K/R), dimethylation (+28.03 Da, K/R), trimethylation (+42.05 Da, K), and crotonylation (+68.03 Da, K). Protein-protein crosslinks were visualized in xiVIEW [24], and conserved domains were annotated using NCBI CD-Search [25].

## 3 Results

### 3.1 Quality of Nuclear Enrichment and Proteomic Data

To assess dataset quality, we combined orthogonal evidence for enrichment of intact nuclei, robust LC–MS/MS performance and identification quality, and biological coherence of the derived protein identifications and interaction resources.

The efficiency of nuclear enrichment was evaluated by fluorescence microscopy of DAPI-stained fractions after sucrose-gradient ultracentrifugation and by comparative SDS-PAGE profiles of whole-cell versus nuclear extracts (Figure 2). Microscopy analysis showed abundant intact nuclei with no or minimal visible contamination (Figure 2a). Coomassie-stained SDS–PAGE revealed a simplified protein profile for nuclear extracts relative to the whole-cell lysate, with prominent low-molecular-weight bands in the expected histone mass range and only subtle DSSO-dependent changes across crosslinker concentrations (Figure 2b) — consistent with proteomewide DSSO crosslinking in complex samples, where crosslinks occur at low stoichiometry and frequently include intramolecular links [10]. Accordingly, DSSO reactivity was primarily validated at the peptide level by identification of crosslinked peptides in the XL-MS analysis (Table S3 and S4), confirming successful *in situ* crosslinking despite the limited sensitivity of gel-based readouts. Altogether, these results support efficient nuclei enrichment and effective yet inherently sub-stoichiometric *in situ* crosslinking within the complex nuclear environment.

**Figure 2.**
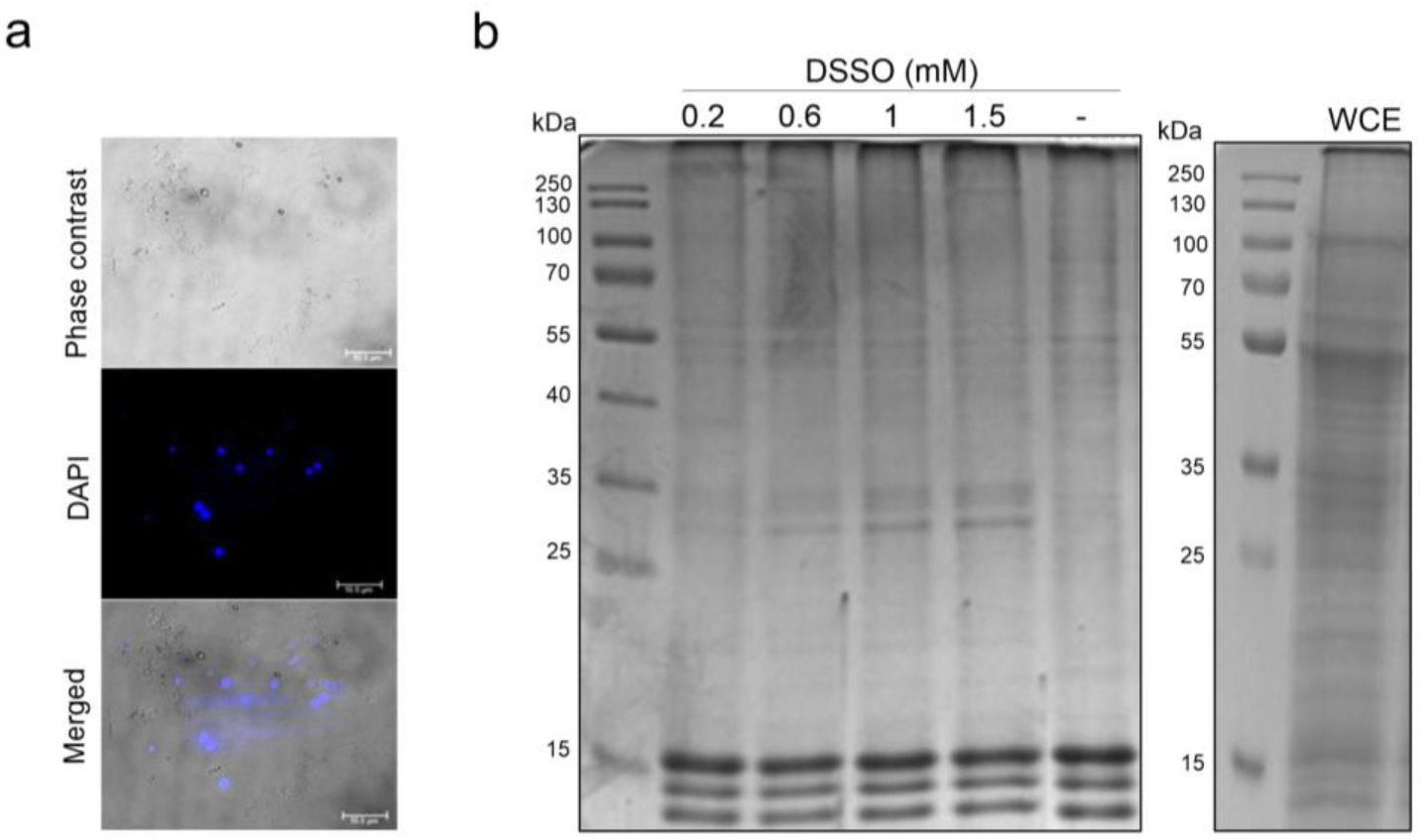
Quality control of the enriched *T. cruzi* nuclear fraction and crosslinking reaction. (a) Representative images of purified nuclei after sucrose-gradient ultracentrifugation, shown as phase-contrast, DAPI fluorescence and merged views (scale bar, 10 µm). (b) Coomassie-stained SDS-PAGE of nuclear extracts prepared from intact nuclei incubated with increasing DSSO concentrations (0.2-1.5 mM), and not incubated (“-”), alongside with a whole-cell extract (WCE) of *T. cruzi* epimastigotes (not incubated with DSSO) also resolved by SDS-PAGE and stained with Coomassie for comparison with the epimastigote nuclear extract on the left. Both gels have acrylamide concentrations of 15%. Molecular weight markers are shown for reference, being the Thermo Scientific PageRuler Prestained Protein Ladder (gel on the left) and Thermo Scientific PageRuler Plus Prestained Protein Ladder (gel on the right).

For the nuclear proteome, peptide and protein identifications were controlled at 1% FDR using a target–decoy strategy implemented in Percolator following Sequest HT and MS Amanda searches, increasing identification robustness and sensitivity for high-resolution Orbitrap MS/MS data, as previously demonstrated [26]. For XL-MS, FDR was controlled at 1% both at the spectral level and at the CSM level in XlinkX [23]. A total of 95,647 PSMs led to the identification of 8,536 peptide sequences belonging to 1,734 proteins (Table S2). On average, proteins were supported by ∼55 PSMs and ∼4 unique peptides. Among the identified proteins, 1,186 were supported by two or more unique peptides, with an average of ∼72 PSMs and ∼6 unique peptides per protein. The remaining 548 proteins supported by a single unique peptide, still exhibited an average of ∼19 PSMs per peptide/protein.

We assessed LC–MS/MS performance and peptide identification quality using standard metrics calculated on the combined identifications obtained from searches performed with and without variable PTMs (Figure 3). Protein identifications were highly concordant between strategies, with 1,660 proteins shared (96%) and only a small number unique to either the proteome-only (55) or PTM-enabled (19) searches (Figure 3a). The missed-cleavage distribution was dominated by fully tryptic peptides (7,502; 89%) and peptides with a single missed cleavage (892; 10%), with only a minor fraction showing ≥2 missed cleavages (52; 1%), consistent with efficient digestion (Figure 3b). In addition, precursor mass errors were tightly centered around zero, with 96.4% of PSMs within ±5 ppm and 83.4% within ±2.5 ppm, indicating stable mass calibration across runs (Figure 3c).

**Figure 3.**
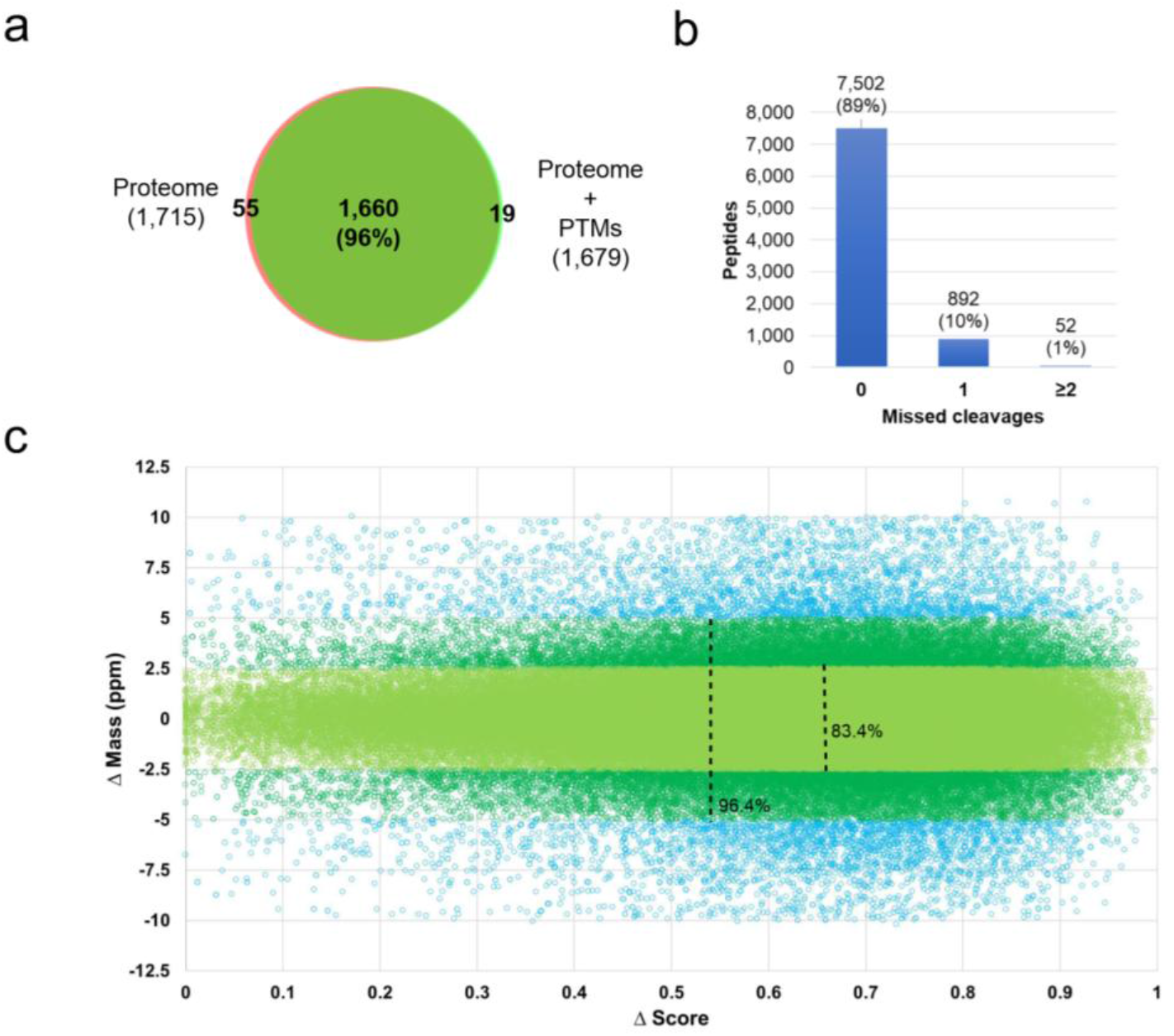
Identification quality metrics for the *T. cruzi* nuclear proteome dataset. (a) Overlap of protein identifications obtained from database searches performed with or without variable PTM settings. (b) Distribution of missed cleavages for peptides in the combined identifications from searches run with and without PTMs. (c) Precursor mass accuracy (ΔMass, ppm) plotted against identification score (ΔScore) for the summed PSM identifications from searches run with and without PTMs.

### 3.2 *T. cruzi* nuclear proteome profiling

Proteome profiling identified 1,734 proteins in total, including 316 proteins identified with PTM-containing peptides (Table S2). Downstream analyses with DeepLoc localization/nuclear-targeting features and GO enrichment consistently indicate predominant nuclear signatures (Figure 4). Based on localization prediction, 1,128 (∼65%) proteins were associated with the nucleus, most of which annotated as nuclear (n = 912, ∼53%) and a subset assigned to the cytoplasm (n = 216, ∼13%) but containing canonical nuclear localization signals (NLSs) and/or nuclear export signals (NESs), suggesting a dynamic nucleocytoplasmic distribution (Figure 4a and Table S5). Notably, the PTM-bearing subset of proteins showed a stronger nuclear bias, with ∼73% assigned to the nucleus. Consistently, Gene Ontology (GO) enrichment across cellular components, biological process, and molecular function highlighted predominantly nuclear terms (Figure 4b). With respect to predicted solubility, the nuclear-targeted subset included both soluble and membrane-associated proteins, with soluble proteins largely predominating (Figure 4c, Table S5). Most membrane-associated proteins were assigned to both soluble and membrane categories, whereas a small subset was classified as membrane-only. Also, most membrane-associated proteins were classified as peripheral membrane proteins.

**Figure 4.**
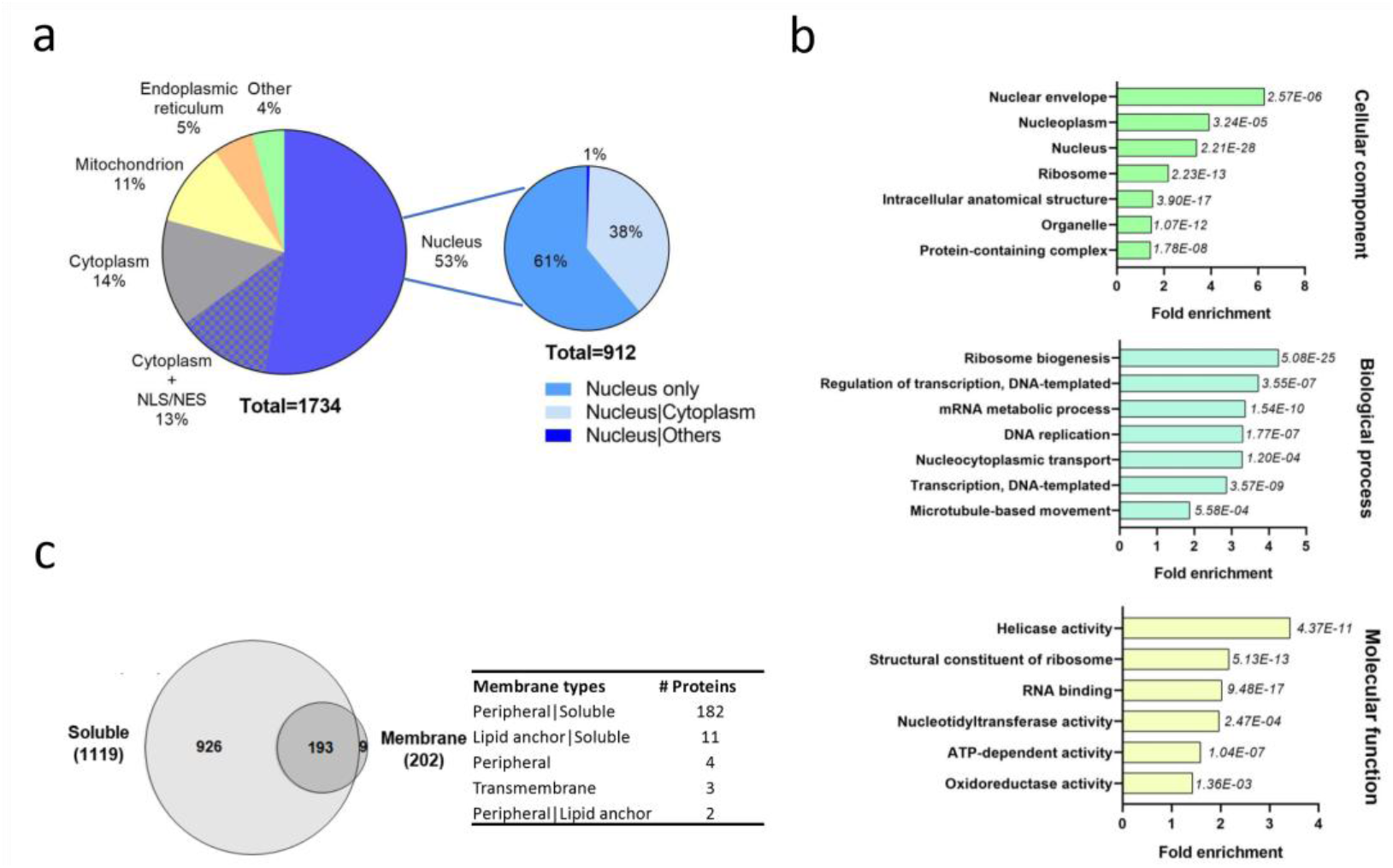
Subcellular localization, solubility/membrane association, and functional enrichment of proteins identified in the *T. cruzi* nuclear fraction. (a) DeepLoc-predicted subcellular localization [17] for all proteins identified (n = 1,734) in the nuclear fraction. A total of 912 proteins (53%) were directly predicted as nuclear, including 557 proteins predicted exclusively as nuclear. For downstream analyses, a broader nuclear-targeted subset (n = 1,128) was defined by combining proteins predicted as nuclear with cytoplasmic proteins carrying predicted NLSs and/or NESs. (b) GO-slim enrichment for all proteins identified (n = 1,734) in the nuclear fraction performed using the TritrypDB GO search tool [16]. Bar plots show fold enrichment for the most enriched terms within Cellular Component, Biological Process, and Molecular Function; significance was assessed using a nominal p-value cutoff of 0.05 and multiple-testing correction (Benjamini–Hochberg and Bonferroni). c) DeepLoc-derived solubility and membrane-association predictions [17] for the nuclear-targeted proteins (n = 1,128; proteins assigned as nuclear plus cytoplasmic proteins carrying predicted NLS and/or NES. Overlap between soluble and membrane-associated assignments is shown, and the inset table summarizes predicted membrane subtypes.

### 3.3 *T. cruzi* protein interaction landscape

The presented nuclear interaction resource integrates two complementary approaches: a STRINGDB-derived network of interactions predicted *in silico* from proteins identified in enriched samples of *T. cruzi* nuclei (Figures 5 and 6); and an exploratory XL-MS–derived set of nuclear protein-protein interactions and associated PTMs identified *in situ* through the chemical crosslinking of enriched samples of *T. cruzi* nuclei (Figure 7).

**Figure 5.**
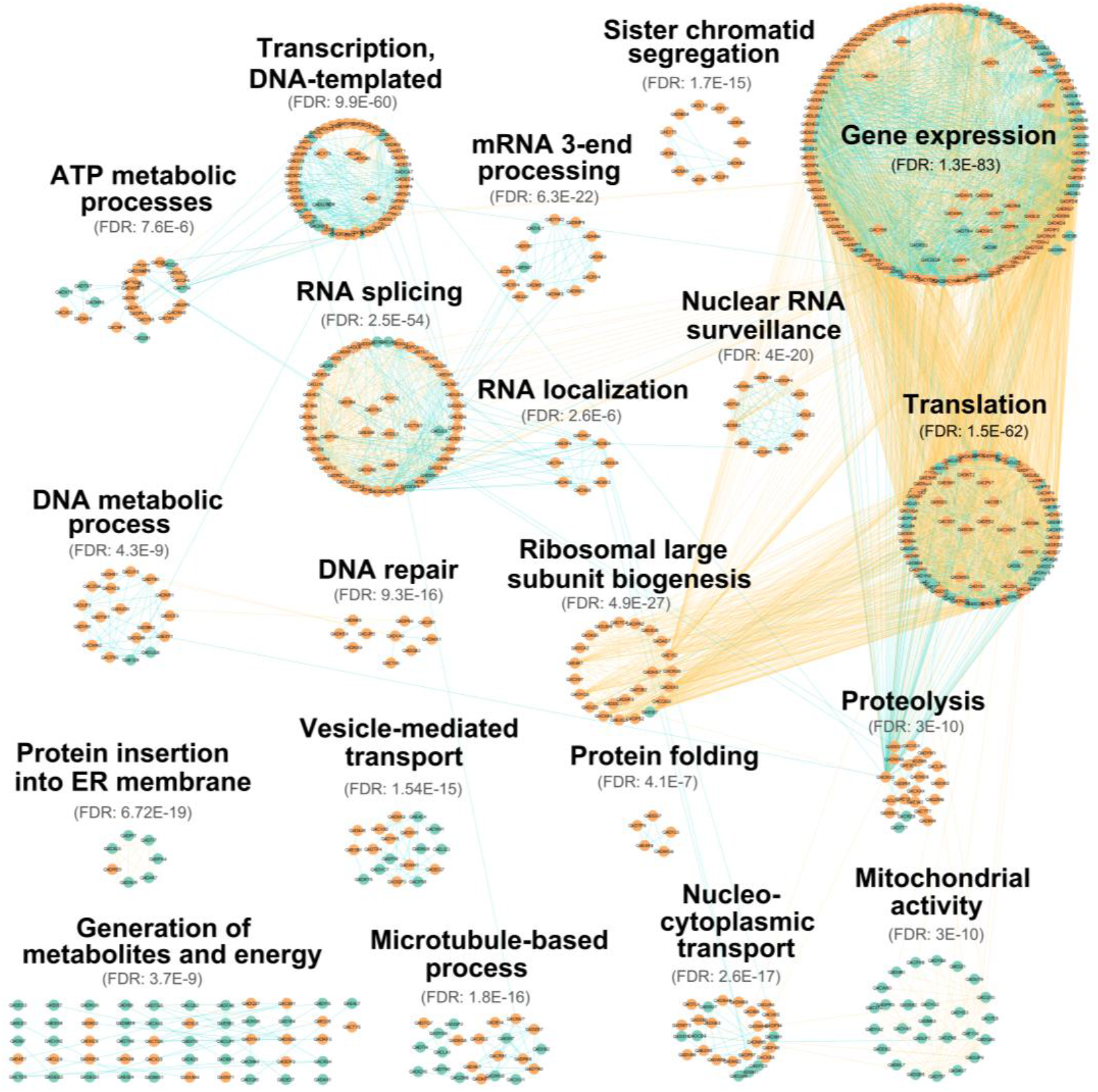
STRINGDB-derived *in silico* interaction map of the *T. cruzi* nuclear proteome. Overview of the MCL [22] clusters derived from the predicted STRINGDB network using the IDs of proteins identified in the *T. cruzi* nucleus as input in StringApp [20] (full STRINGDB mode; minimum required interaction score = 0.7; inflation = 4). Each mini-network represents one MCL cluster; clusters with significant functional enrichment are labeled with the top enriched GO Biological Process term and its FDR. Node colors indicate proteins predicted by DeepLoc [17] as nuclear (orange) versus other localizations (teal). For visualization, displayed edges were restricted to a high-confidence subset (score ≥ 0.9) supported by experimental evidence (orange) or curated database evidence (blue) available within the STRING database [19].

**Figure 6.**
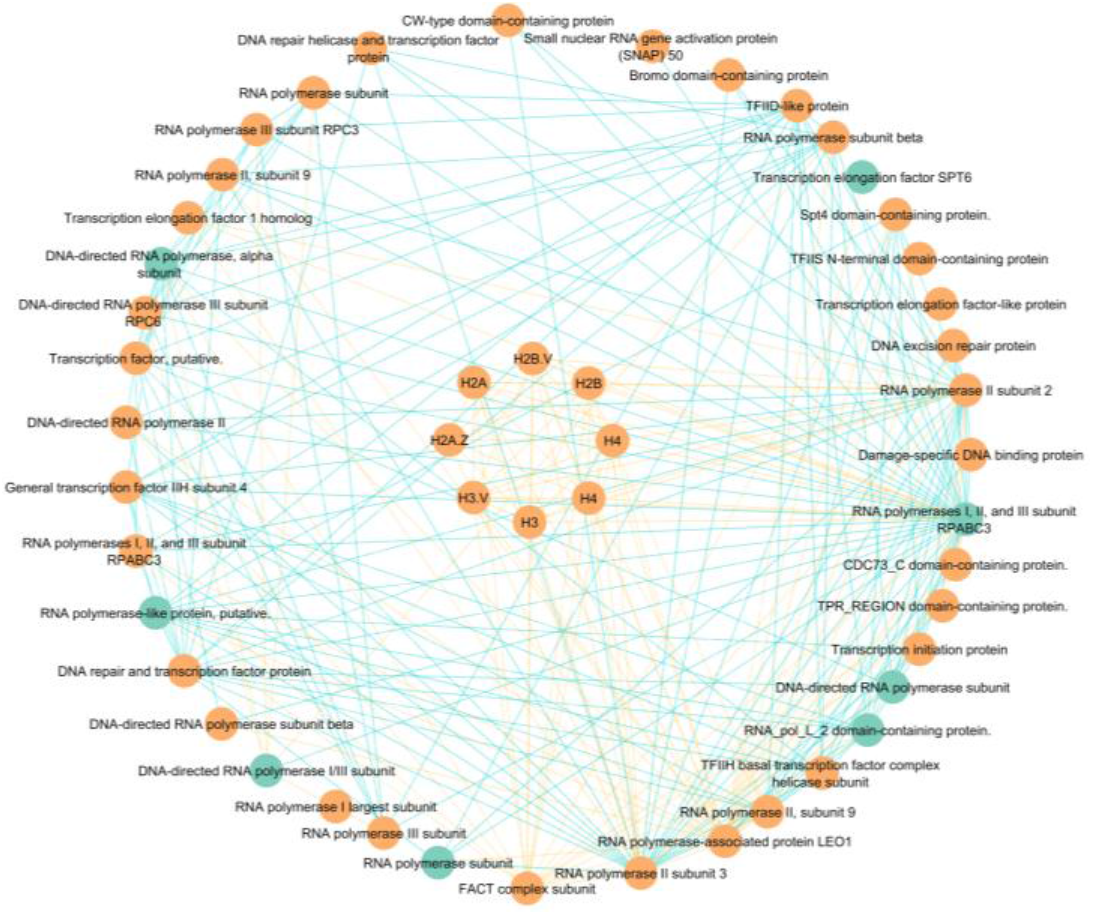
STRINGDB-derived *in silico* histone-centered interaction subnetwork of the *T. cruzi* nuclear proteome. The MCL [22] cluster annotated as “transcription, DNA-templated” from the STRINGDB-derived interaction network is shown, highlighting canonical (H2A, H2B, H3, and H4) and variant (H2A.Z, H2B.V, and H3.V) histones and their firstand second-degree interaction partners. Node colors indicate proteins predicted by DeepLoc [17] as nuclear (orange) versus other localizations (teal). Edges shown correspond to the high-confidence visualization subset (score ≥ 0.9) and are restricted to interactions supported by experimental evidence (orange) or curated database evidence (blue) available within the STRING database [19].

**Figure 7.**
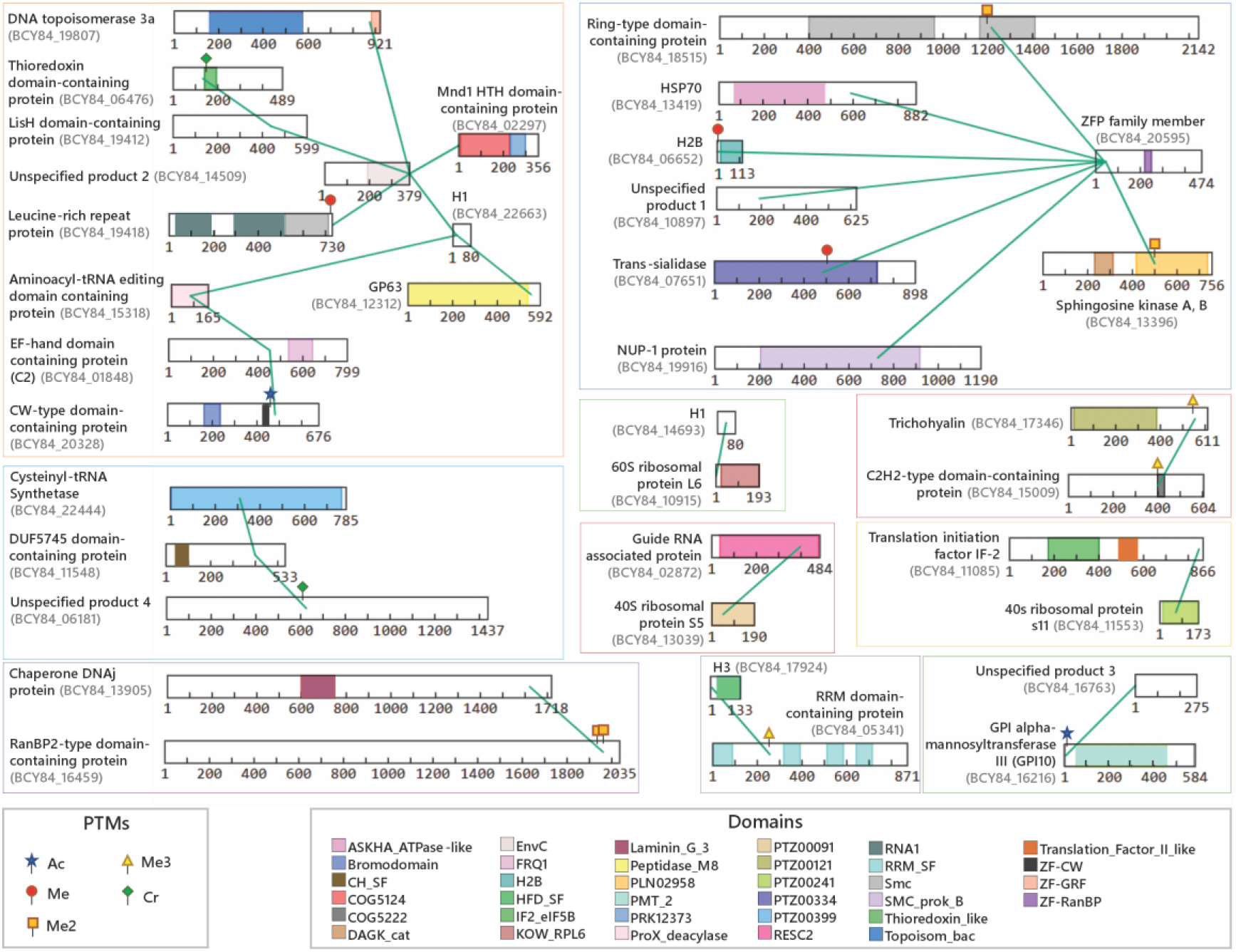
Exploratory XL-MS interaction network derived from crosslinked peptides in the nuclear-enriched fraction of *T. cruzi*. Protein-protein crosslinks were visualized in xiVIEW [24]. Each protein is displayed as a linear bar scaled by residue number and identified by its protein ID in the TriTrypDB database [16] v.68. Proteins with no characterized function and/or annotation in *T. cruzi* or among their orthologues in TriTrypDB/Uniprot were designated as “Unspecified product”. Green connectors denote inter-protein crosslinks, and connector endpoints indicate the linked residue positions. Conserved domains were annotated using NCBI CD-Search [25] and are shown as colored blocks along each sequence (domain color key inset). The colored rectangles separate different clusters. PTMs detected on crosslinked peptides are marked at their corresponding sites (PTM key inset): acetylation (Ac), monomethylation (Me), dimethylation (Me2), trimethylation (Me3), and crotonylation (Cr).

The *in silico* interaction network organizes a total of 608 proteins detected in the nuclear preparation into 20 interaction clusters with coherent functional signatures, annotated using the top enriched GO Biological Process terms (Figure 5; Tables S6 and S7). The most prominent clusters are dominated by nuclear-related processes, such as gene expression, DNA-templated transcription, RNA splicing, mRNA 3′-end processing, nuclear RNA surveillance, DNA repair, and ribosome biogenesis, consistent with the nuclear-enriched character of the dataset. Proteins predicted as nuclear are prevalent within these modules, whereas proteins assigned to other localizations are more frequent in clusters linked to processes such as mitochondrial activity or ER-associated pathways. Within this network, the cluster annotated as “transcription, DNA-templated” corresponds to a histone-centered interaction neighborhood (Figure 6), highlighting canonical (H1, H2A, H2B, H3, and two H4 isoforms) and variant histones (H2A.Z, H2B.V, and H3.V) associated with 41 interaction partners, 33 of which displayed nuclear localization, including transcription-associated factors and RNA polymerase subunits. These findings provide a focused resource of candidate histone-associated proteins within the *T. cruzi* nuclear proteome.

In complement to the *in silico* network, we generated an exploratory XL-MS dataset from the enriched intact nuclei, capturing native protein-protein interactions for 36 proteins (Figure 7; Tables S3 and S4). Most crosslinked proteins were assigned to the nucleus, and the interaction map includes, among others, histones (H1, H2B, and H3) and proteins associated with RNA-related processes and translation. In addition, PTMs annotated on crosslinked peptides—primarily lysine acylation and methylation states, as well as crotonylation—were observed at or near linked sites, providing an integrated view of candidate interaction partners and nearby modification sites for downstream exploration. Four of the crosslinked proteins are still annotated as “unspecified product” in TriTrypDB/UniProt; for these, conserved-domain predictions highlighted RNA-binding and other functional features, providing additional functional context and supporting hypothesis generation. Thus, this XL-MS resource complements the STRINGDB-derived network by providing an experimentally supported subset of nuclear interaction partners with associated site-proximal PTM annotations.

## 4 Discussion

Compared with the previously published *T. cruzi* nuclear proteome [9], the present dataset expands by approximately 2-fold the number of described nuclear proteins, including a larger number of proteins with predicted nuclear localization. Differences between the two datasets may reflect multiple factors, including biological variation, sample preparation procedures, fractionation strategies, mass spectrometry instrumentation, database versions, and data-processing workflows. Notably, the workflow employed in the present study included DSSO crosslinking, SCX fractionation, and analysis using a high-resolution Orbitrap Fusion Lumos instrument. However, the relative contribution of these factors to the increased proteome coverage cannot be determined from the available data. Therefore, the two studies should be regarded as complementary resources that collectively expand the current coverage of the *T. cruzi* nuclear proteome.

This dataset supports investigations of nuclear composition, organization, and function in the *T. cruzi* parasite. Potential downstream applications include: (i) comparative nuclear proteomics, to identify conserved and parasite-specific nuclear components; (ii) nuclear, chromatin and histone interaction studies, to select candidate nuclear protein complexes, histone ligands and chromatin-associated proteins that can be prioritized for functional analysis using genetics, ChIP-based approaches or biochemical assays; and (iii) PTM-centered analyses, in which modified sites detected on histones and non-histone proteins — particularly those located near exploratory crosslinks — can be integrated with independent PTM datasets to assess how specific modifications modulate nuclear interactions. In addition, the present resource may be integrated with previously published *T. cruzi* nuclear proteome [9] and chromatin-associated proteome [27] studies to support comparative and integrative analyses of nuclear organization, chromatin regulation, and nuclear protein composition in this parasite.

For reuse, users should consider the confidence information provided in the annotation tables (e.g., FDR-controlled identifications, peptide evidence/uniqueness, STRINGDB confidence scores and evidence channels, and XL-MS confidence metrics) and apply filtering criteria appropriate to the intended application. Proteins identified based on a single unique peptide (n = 548) that passed the stringent validation criteria applied during data processing, including Percolator-based validation and a strict 1% false discovery rate (FDR) at the PSM, peptide, and protein levels, were retained in the dataset. Most of these proteins were additionally supported by multiple PSMs assigned to the same unique peptide. While proteins identified by multiple unique peptides generally provide greater confidence, single-peptide identifications may include biologically relevant low-abundance proteins and were therefore not excluded from the resource. Users should evaluate this subset according to the requirements of their specific applications. For more conservative downstream analyses, additional filters such as requiring at least two unique peptides may be applied.

## 5 Limitations

There are some possible biological and methodological reasons for the presence of nonnuclear proteins in our dataset. Even highly enriched nuclear preparations may retain proteins associated with cellular structures that physically or functionally interact with nuclei, including mitochondria, endoplasmic reticulum, ribosomes, cytoskeletal components, and membrane-associated complexes. Consequently, a proportion of proteins assigned to non-nuclear compartments may reflect co-purification or carryover inherent to biochemical fractionation procedures rather than purely technical contamination [28, 29]. This consideration may be particularly relevant in *T. cruzi* and other trypanosomatids, whose organelles are closely associated within a highly organized microtubule network and exhibit extensive structural and functional interactions. Such features have previously been noted as factors influencing the interpretation of subcellular fractionation datasets in these organisms [9, 30]. In addition, some proteins may exhibit dual localization or dynamic nucleo-cytoplasmic trafficking. Finally, subcellular localization assignments in our study were based on DeepLoc predictions, and prediction-based approaches may have reduced accuracy when applied to evolutionarily divergent non-model organisms such as *T. cruzi*. Therefore, proteins assigned to non-nuclear compartments should not be interpreted solely as contaminants but rather as a heterogeneous group that may include carryover proteins, dual-localized proteins, proteins associated with nuclear-adjacent cellular structures, and proteins undergoing nucleo-cytoplasmic shuttling.

Protein identifications reported in this study were generated using the *Trypanosoma cruzi* Dm28c 2017 protein database available in TriTrypDB release 68. Future updates to TriTrypDB may refine gene models, protein annotations, and functional assignments, potentially affecting downstream annotation and biological interpretation.

For the *in silico* interactome, we reduced spurious connections by focusing on high- and very-high-confidence STRINGDB interactions (minimum scores of 0.7 and 0.9, respectively). Community detection organized the nuclear proteome into functionally coherent clusters, enriched for canonical nuclear processes (Figures 3–4), supporting the biological plausibility of the inferred network. For reuse, we provide the full interaction tables retrieved at score ≥0.7, whereas the figure panels show a stringent, high-confidence subset (score ≥0.9) restricted to experimentally supported or curated database edges, which reduces visual complexity while preserving interpretable modules. Overall, the combination of high STRINGDB confidence, recovery of established nuclear assemblies, and coherent functional clustering supports a robust interaction framework anchored in the experimental proteome.

In parallel, XL-MS data are presented as an exploratory, orthogonal layer that complements the STRINGDB-derived network, in XlinkX under stringent FDR control (1% at both spectral and CSM levels; Tables S3 and S4). The resulting crosslinks involve histones (H1, H2B, H3) and additional proteins linked to RNA-related processes, translation, and uncharacterized nuclear factors. Although crosslink coverage is modest — consistent with the challenges of deep XL-MS in complex nuclear samples and the use of a preconfigured DSSO acquisition scheme — the individual identifications are supported by nuclear localization prediction and network context. Several crosslinked peptides carry PTMs (acetylation, methylation, crotonylation) at or near the crosslinked residues, supporting the biological relevance of these sites. Accordingly, the XL-MS dataset is not intended as a comprehensive nuclear interactome, but as a set of site-resolved hypotheses on specific contacts and potentially PTM-modulated interfaces that can be revisited as XL-MS strategies continue to improve.

## 6 Conclusions

Overall, the data and associated annotation tables are suitable for reuse in multiple contexts, including comparative nuclear proteomics across trypanosomatids, integration with transcriptomic and chromatin datasets, and prioritization of candidates for follow-up and functional studies.

## Supplementary Materials

Table S1. Relationship between deposited RAW files, experimental metadata, analysis workflows, and output files. This table provides a comprehensive overview of all deposited mass spectrometry data files within PRIDE, including RAW file identifiers, PRIDE file type, corresponding experimental conditions, and search type. The table serves as a reference for tracing each dataset from acquisition through computational analysis and facilitates reproducibility and data reuse.

Table S2. Protein identifications from LC–MS/MS searches. Protein identifications exported from Proteome Discoverer, including protein identifiers and annotations with peptide evidence and identification confidence information as reported by the software. The column “Search type” indicates the database search workflow(s) in which each protein was identified: “Proteome” denotes proteins identified exclusively in the standard proteome search; “PTMs” denotes proteins identified exclusively in the PTM-enabled search; and “Proteome-PTMs” denotes proteins identified in both search workflows. Both searches were performed using the same LC–MS/MS RAW files but differed in the set of variable modifications included during database searching, as described in the Methods section (“Proteome and PTM searches”). Where applicable, PTM annotations associated with identified peptides are included.

Table S3. Summary of crosslinks identified in the XL-MS dataset. Crosslink-level output from XlinkX, summarizing identified inter- and intra-protein links with the corresponding linked proteins/sites and associated peptide-level annotations and confidence information as reported by the software.

Table S4. Crosslink spectrum matches supporting identified crosslinked peptide pairs. CSM-level output from XlinkX, providing acquisition identifiers and confidence/scoring information together with fragment-ion evidence fields (as reported by the software), enabling inspection and re-evaluation of individual crosslink identifications.

Table S5. Subcellular annotation and solubility/membrane predictions for identified proteins. DeepLoc-derived annotations for each identified protein, including subcellular compartment assignment, nuclear-targeting features (predicted NLS/NES), and solubility/membrane-association outputs, including predicted membrane subtypes when available.

Table S6. Node table for the STRINGDB interaction network and community assignments. Node table exported from STRINGDB via stringApp/Cytoscape for the released network, including protein identifiers and network attributes as provided by STRINGDB/Cytoscape, together with Markov Cluster Algorithm (MCL) cluster assignments used for module/community analyses.

Table S7. Edge table for the STRINGDB interaction network used for clustering and downstream filtering. Edge table exported from STRINGDB via stringApp/Cytoscape for the released network, listing interacting protein pairs with STRINGDB interaction scores and evidence-channel metadata (as provided by STRINGDB), supporting MCL-based module analyses and downstream visualization filtering.

## Author Contributions

For research articles with several authors, a short paragraph specifying their individual contributions must be provided. The following statements should be used “Conceptualization, L.M.F.G. and R.F.A; methodology, L.M.F.G., M.F. and R.F.A.; validation, R.F.A. and L.M.F.G.; formal analysis, R.F.A. and L.M.F.G.; investigation, M.F. and R.F.A.; resources, L.M.F.G.; data curation, R.F.A.; writing—original draft preparation, R.F.A; writing—review and editing, R.F.A and L.M.F.G.; visualization, R.F.A and L.M.F.G.; supervision, L.M.F.G.; project administration, L.M.F.G.; funding acquisition, L.M.F.G. All authors have read and agreed to the published version of the manuscript.

## Data Availability Statement

The original data presented in the study are openly available in ProteomeXchange Consortium via the PRIDE [31] partner repository with the dataset identifier PXD071382 (available at https://doi.org/10.6019/PXD071382). The PRIDE submission includes: 16 RAW files (.raw), corresponding to the vendor files as acquired; 16 Mascot Generic Format files (.mgf), corresponding to MS/MS peak lists exported from Proteome Discoverer; three mzML files (.mzML) and three mzIdentML results files (.mzid), corresponding to peptide/protein, PTMs, and crosslink identifications from Proteome Discoverer searches. Table S1 describes all deposited PRIDE files and their relationship to the corresponding experimental conditions, LC–MS/MS runs, and downstream search workflows.

## Acknowledgments

The authors thank CNPq, CAPES and Fiocruz for financial support. They are also grateful to the Instituto Carlos Chagas Mass Spectrometry Facility (RPT02H), Technological Platforms Network, Oswaldo Cruz Foundation (FIOCRUZ), for access to instrumentation and infrastructure.

## Abbreviations

Ac: Acetylation
AGC: Automatic Gain Control
AmAc: Ammonium acetate
CID: Collision-Induced Dissociation
CSM: Crosslink Spectrum Match
DAPI: 4’,6-Diamidino-2-phenylindole
DDA: Data-Dependent Acquisition
DSSO: Disuccinimidyl Sulfoxide
ΔMass: Precursor mass error
ΔScore: Identification score difference
EDTA: Ethylenediaminetetraacetic acid
ER: Endoplasmic Reticulum
ETD: Electron Transfer Dissociation
FDR: False Discovery Rate
FTMS: Fourier Transform Mass Spectrometry
GO: Gene Ontology
hPTMs: Histone Post-Translational Modifications
ITMS: Ion Trap Mass Spectrometry
LC–MS/MS: Liquid Chromatography coupled to Tandem Mass Spectrometry
LIT: Liver Infusion Tryptose
MCL: Markov Cluster Algorithm
Me: Monomethylation
Me2: Dimethylation
Me3: Trimethylation
mgf: Mascot Generic Format
m/z: Mass-to-charge ratio
NES: Nuclear Export Signal
NLS: Nuclear Localization Signal
PPI: Protein–Protein Interaction
ppm: Parts per million
PSM: Peptide-Spectrum Match
PTM: Post-Translational Modification
ptmRS: PTM site localization algorithm
RSLC: Rapid Separation Liquid Chromatography
SCX: Strong Cation Exchange
SDS: Sodium Dodecyl Sulfate
SDS-PAGE: Sodium Dodecyl Sulfate Polyacrylamide Gel Electrophoresis
STRING: Search Tool for the Retrieval of Interacting Genes/Proteins
Tris-HCl: Tris(hydroxymethyl)aminomethane hydrochloride
WCE: Whole-Cell Extract
XL-MS: Crosslinking Mass Spectrometry

